# Rett syndrome – biological pathways leading from MECP2 to disorder phenotypes

**DOI:** 10.1101/062786

**Authors:** Friederike Ehrhart, Susan L.M. Coort, Elisa Cirillo, Eric Smeets, Chris T. Evelo, Leopold M. Curfs

## Abstract

Rett syndrome (RTT) is a rare disease but still one of the most abundant causes for intellectual disability in females. Typical symptoms are onset at month 6-18 after normal pre-and postnatal development, loss of acquired skills and severe intellectual disability. The type and severity of symptoms are individually highly different. A single mutation in one gene, coding for methyl-CpG-binding protein 2 (MECP2), is responsible for the disease. The most important action of MECP2 is regulating epigenetic imprinting and chromatin condensation, but MECP2 influences many different biological pathways on multiple levels. In this review the known changes in metabolite levels, gene expression and biological pathways in RTT are summarized. It is discussed how they are leading to some characteristic RTT phenotypes and identifies the gaps of knowledge, namely which phenotypes have currently no mechanistic explanation leading back to MECP2 related pathways. As a result of this review the visualization of the biologic pathways showing MECP2 up-and downstream regulation was developed and published on <u>WikiPathways</u>.

## 1 Introduction

Rett syndrome (RTT; <u>MIM:312750</u>) occurs in 1:10.000 girls at the age of 12 (1). It is considered a rare disease since it affects fewer than 1 in 2000 individuals (2), but it is still one of the most abundant causes for intellectual disability in females. RTT was first described in 1966 by the Viennese pediatric Andreas Rett, who observed the typical hand movements (“hand washing”) of his patients (3, 4). Cause of RTT is in most cases a *de novo* mutation of <u>*MECP2*</u> (methyl-CpG-binding protein 2) gene; which was discovered by Amir et al. (5). However, as stated by Neul et al., “not all mutations in *MECP2* cause RTT and not all RTT patients have mutated *MECP2”*. Mutations in two other genes can also cause a RTT like phenotype, i.e. *FOXG1* and *CDKL5,* this was formerly considered as RTT but is now defined as RTT like syndrome (6).

RTT was considered a neurodevelopmental disorder but since some of the main symptoms were found to be reversible (7) researchers and clinicians tend to categorize it as a neurological disorder now (8). RTT was classified as an autism spectrum disorder before but not anymore. Patients often develop autistic features like social withdrawal but only during a certain stage of development (9). Although RTT has some autistic features/phases these usually disappear with time and adult RTT females are quite socially active again (10). *MECP2* mutations are rarely found in autism patients and if, they are termed “Autism with MECP2 mutation” (6).

In this review, we summarize the knowledge about the *MECP2* gene and protein and discuss how they cause RTT phenotypes. In addition, the current insights in the downstream regulation activity of MECP2 in RTT females will be discussed. This review integrates database knowledge from <u>Ensembl, OMIM, UniProt, The Human Protein Atlas</u>, and <u>Gene Ontology</u> and a biologic pathway was developed and published on <u>WikiPathways</u> do visualize the mechanistic action of MECP2.

## 2 Rett syndrome phenotype and diagnosis

Clinical diagnosis of RTT requires the following symptoms: decreased, arrested and retarded development of motoric and communication skills after 6-18 months of normal postnatal development, development of stereotypic movements and loss of purposeful movement. Although the onset of typical disorder symptoms after the age of 6 months is characteristic for RTT, observations of parents that “something is wrong with this child”, are often made before. RTT females are typically severely intellectual disabled, suffer under microcephaly and seizures. Additionally, they often develop symptoms like cardiac and breathing abnormalities, low muscle tension, autistic like behavior, scoliosis (and other osteopathies), sleeping problems and hormone disequilibrium (summary in Table 1]). Typical noticeable laboratory results are EEG (and EKG) abnormalities, atypical brain glycolipids, altered neurotransmitter, creatine and growth factor levels, and alkalosis. Most of the symptoms can be related to disturbed neuronal function but some of them are caused or influenced by alterations which are not yet elucidated (4, 11, 12). It is also unknown whether it is the often in RTT observed changes of carbondioxide metabolism that cause respiratory problems or that dysfunctional brain stem neurons are responsible for breathing abnormality (4).

**Table 1:** Diagnosis criteria and characteristic symptoms of RTT [* Source: <u>OMIM#312750</u>]

|  |  |
| --- | --- |
| **Diagnosis criteria** | • Arrested development between 6-18 months<br>• Regression and loss of acquired skills<br>• Loss of speech<br>• Stereotypic movements<br>• Gait abnormalities |
| **Typical additional symptoms** | • Cardiac abnormalities<br>• Low muscle tension/hypotonia/ dystonia causing e.g. incontinence, constipation, chorea, bruxism, ataxia, spasticity<br>• Impaired, autistic like social behavior: avoidance of eye contact, lack of social/emotional reciprocity, impaired use of nonverbal behavior, screaming fits, inconsolable crying<br>• Breathing abnormalities<br>• Consequences of mal/undernutrition(?): short stature, osteopathy, vitamin D deficiency, skoliosis<br>• Sleeping problems<br>• Hormone disequilibrium, e.g. thyroid hormones<br>• Microencephaly<br>• Seizures<br>• Intellectual disability |
| **Noticeable laboratory results** | • EEG abnormalities<br>• Atypical brain glycolipids<br>• Elevated CSF levels of beta-endorphin and glutamate<br>• Reduction of substance P<br>• Decreased level of CSF nerve growth factors (BDNF)<br>• Abnormal creatin levels<br>• (Respiratory) alkalosis<br>• Smaller and more densely packed neurons<br>• Shorter, less complex and less dendritic spines |

\*Source: OMIM#312750

## 3 MECP2 gene, transcript and protein

The *MECP2* gene is highly conserved in Euteleostomi (bony vertebrates). The <u>NCBI</u> HomoloGene/UniGene database gives detailed information about gene homologues in 10 mammalian, 2 amphibian and 1 bony fish species (13). The human *MECP2* is located on chromosome X, position 154,021,573-154,137,103 (reverse strand) (according to Ensembl, version 84, genome build GRCh38.p5 (GCA_000001405.20)) and there are currently 21 transcripts known, two of these are protein coding (Figure 1). Due to dosage compensation *MECP2* is inactivated in one X-chromosome in females and the degree of inactivation is assumed to contribute to the difference in phenotypes for RTT (14). RTT is due to its location on the X-chromosome more often observed in females. Hemizygous males with a severe mutation are generally not viable, but there are several non-lethal mutations which can lead to severe congenital encephalopathy, RTT-like syndrome, and mild to severe intellectual disability in males (15). Mosaic expression with only wild-type *MECP2* active in females are possible but supposed to be extremely rare (14).

**Figure 1.**
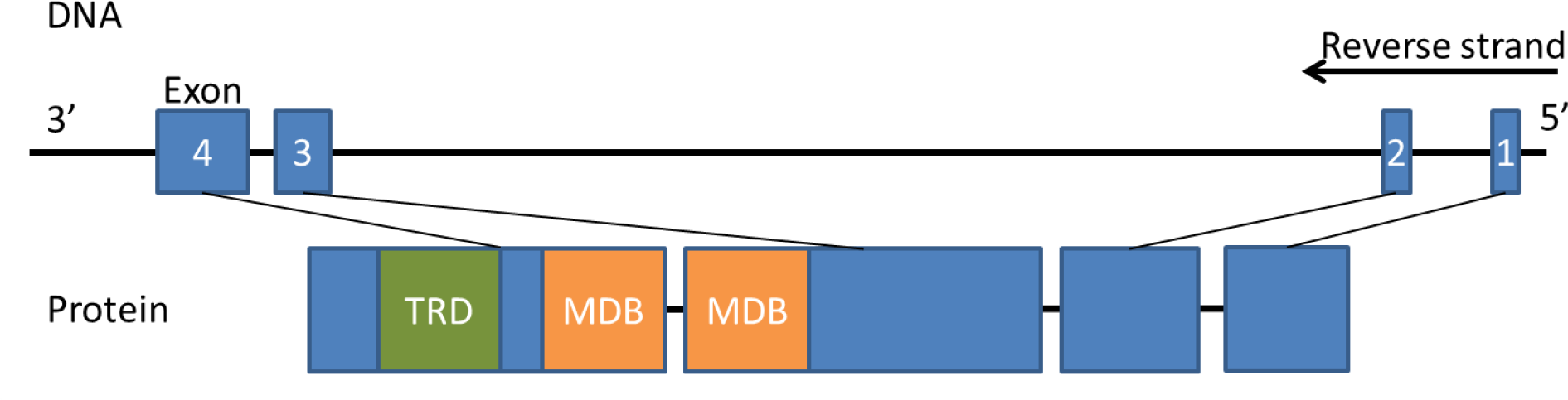
MECP2 gene and protein. *MECP2* gene is about 10505 bp long and has 4 exons. The protein has 486 amino acids and consists of 6 distinct domains whereas the methyl-DNA binding (MDB) and the transcriptional repression domain (TRD) are the most important for function.

The transcription and translation of MECP2 is highly regulated (16) (Table 2 and Figure 2). There are several cis-and trans-regulatory elements for *MECP2* gene expression regulation known. Cis regulatory elements, including promotor elements, are loci on the DNA which act as binding sites for transcription factors and activate or repress gene expression. Trans-elements affect the regulation in an indirect way and can be located close or far away. They can for instance include genes that encode transcription factors for this specific gene. Translation of MECP2 can be regulated by a set of microRNAs (17-25). MicroRNAs are small non-coding RNAs, that repress translation mRNA into protein by binding to the 3′ untranslated region of the mRNA. The regulation of MECP2 expression, stimulation and repression, is visualized in Figure 2 (transcriptional and translational regulation of MECP2) which is derived from <u>WikiPathways</u>.

**Table 2:** Regulation of *MECP2* expression by transcription factors and microRNA

| Type | Transcription factors/microRNA |
| --- | --- |
| Transcription factors targeting *MECP2* cis-elements (17-19) | Activation by BRN3, MYT1, SP1, SP3, C/EBP, CTCF, E2F1, TAF1, TAP1 |
|  | Repression by REST, BRN2, BCL6, |
| Trans-regulatory elements of *MECP2* (20) | Activation by HNRNPF |
|  | Repression by HNRNPH1 |
| miRNA (posttranscriptional repression) (21-25) | hsa-miR-483- 5p, hsa-miR-132-5p, hsa-miR-152-3p, hsa-miR-199a-3p, hsa-miR-30a-3p, and hsa-miR-130b-3p |

**Figure 2.**
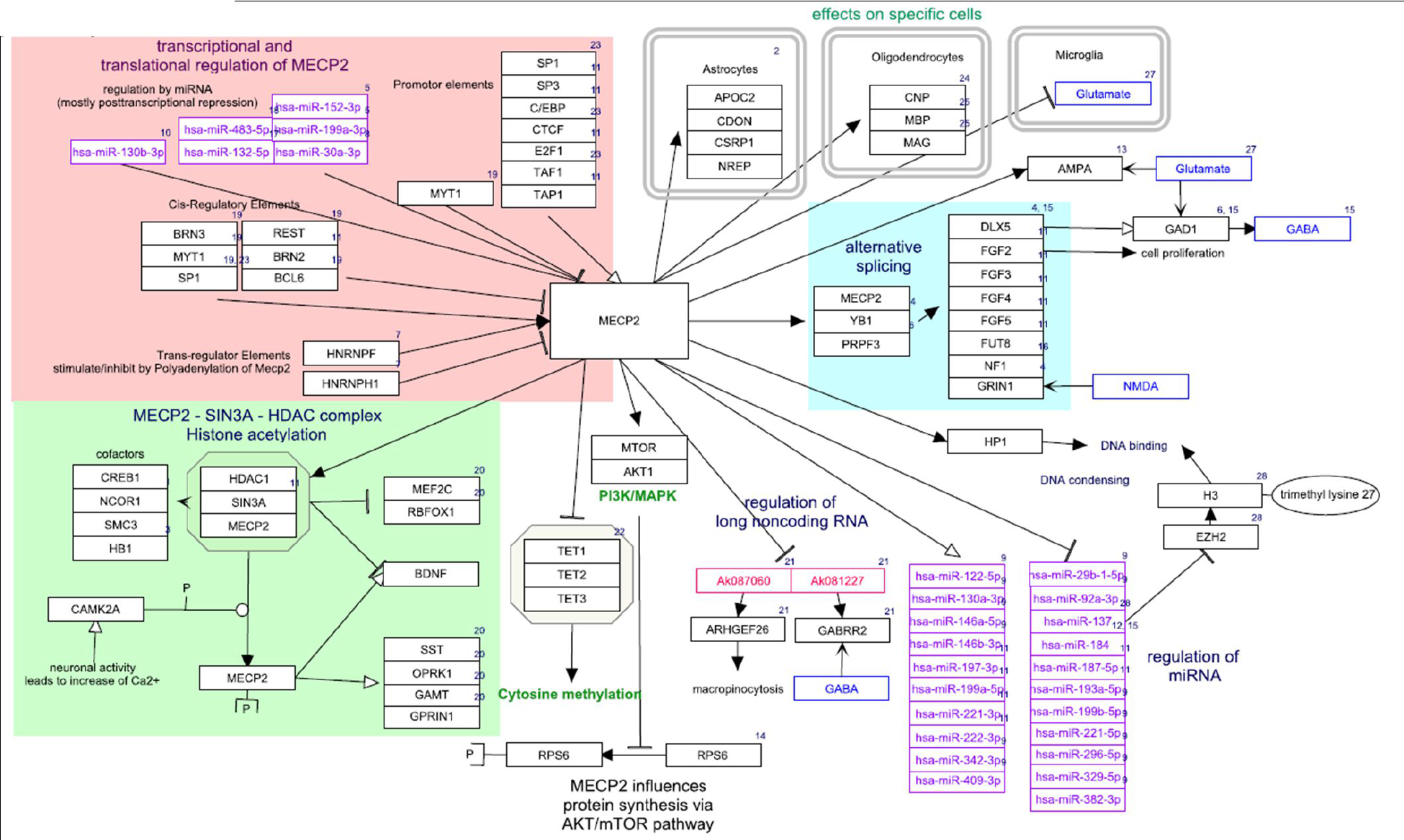
MECP2 pathway from WikiPathways (wikipathways.org/instance/WP3584). Visualization of MECP2 being regulated by several upstream elements, e.g. promotor elements and microRNAs, and MECP2 regulating the expression and splicing of several downstream transcripts, proteins and miRNAs.

The two coding transcripts are isoforms of MECP2, long e1 and short e2, while e1 seems to be the more important one (26, 27). Itoh et al. observed that specific deactivation of e2 did not influence normal neurodevelopment while loss of e1 led to RTT (28). The MECP2 protein has at least six biochemically distinct domains (29). Two of them are most important for the protein function: the (84 amino acids) methyl-CpG binding domain (MDB) which is the one which selectively binds 5MeCyt and the transcriptional repression domain (TRD) (102 amino acids) which binds cofactors attracting histone deacetylase and finally leading to transcription repression as explained in the chapter 4 (Figure 1) (29, 30). Interestingly, MBD is the only structured domain (α-helix) while 60% of MECP2 is unstructured (29). For visualization of the structure see PDB (Protein Databank in Europe) (MECP2 ID: <u>3c2i</u> and <u>1qk9</u>) and the <u>Interpro ID</u> is IPR017353.

MECP2 protein is most abundant in brain but also enriched in lung and spleen tissue (31). However, according to the <u>human protein atlas</u> MECP2 protein (and its transcript) is found in almost every tissue. The cell type with the highest expression level is neurons, but MECP2 expression is also high in glial cells, astrocytes and oligodendrocytes where it influences specific gene expression (see Figure 2, effects on specific cells). In neurons an expression level of about 1.6×10^7^ protein copies per nucleus was estimated by Skene et al. It is about the same number as nucleosomes or 5-methyl-cytosine (5MeCyt) spots on the DNA, leading to the suggestion that every spot might be covered by one MECP2 (32).

## 4 MECP2 function

MECP2 is a multifunctional protein which influences gene expression and metabolism on many levels (8) (Figure 3). The main function of MECP2 is to recognize and bind specifically methylated cytosine residues in the DNA (namely 5MeCyt) that are enriched with A/T bases adjacent (33). MECP2 binds also but with lesser affinity to hydroxymethylated DNA (namely 5-hydroxy methylated cytosine, 5OHMeCyt). Mutations in MECP2, which lead to loss of specific 5MeCyt binding functions are known to cause RTT (34).

This is in line with the Gene Ontology classification for the main molecular functions of MECP2: DNA binding, namely double-stranded methylated DNA and protein binding, namely histone deacetylase. The actual full Gene Ontology annotation of MECP2 can be found online e.g. <u>Ensembl database MECP2 entry</u>.

**Figure 3.**
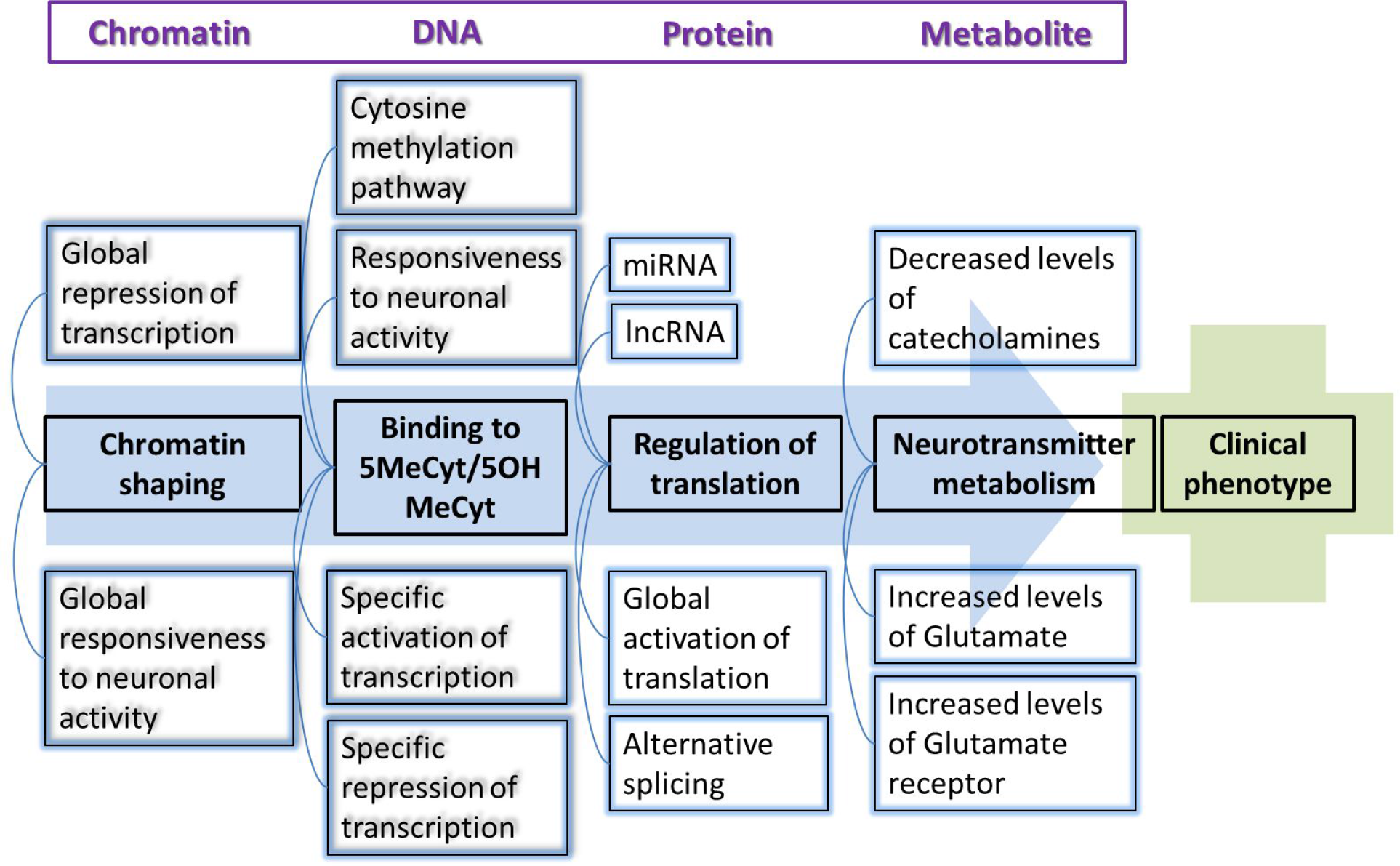
MECP2 and its different levels of influence to chromatin structure, DNA binding, protein and metabolite level leading to clinical phenotypes. 5MeCyt = methylated cytosine, 5OHMeCyt = hydroxylated cytosine.

The molecular functions of MECP2 are known to influence various biological mechanisms, which are summarized and visualized in the pathway Figure 2, namely 1) MECP2 influences global translation by enhancing the AKT/mTOR signaling pathway (35), 2) Alternative splicing of downstream gene products is affected because MECP2 forms a complex with <u>YB1</u>, an important splicing factor (19, 36-40), 3) Expression of various microRNAs and long non-coding RNAs is regulated by MECP2 (20, 45, 47-49), and 4) MECP2 triggers the chromatin compaction at methylated DNA sites which regulates the transcription of adjacent genes (34, 37-41). The last one is an important (and best investigated) pathway and will be explained in detail below.

## 5 MECP2 as epigenetic regulator

Methylation of DNA is part of epigenetic gene expression regulation, where DNA is modified without changing the genetic code. Most transcription factors are unable to bind to methylated DNA so methylation usually silences a gene. Furthermore, methylated DNA is – via MECP2 mediated cofactor binging – a binding site for histone deacetylase (HDAC) which increases DNA compaction by removing certain acetyl residues from lysine at the histone tail allowing them to get closer to each other (Figure 2, MECP2-HDAC complex). So, methylated DNA is tightly wrapped around histone proteins and the access of transcription factors is physically inhibited (30). About 1% of DNA is methylated in humans and the methylation sites are often in regions with a high occurrence of CG, so-called CpG islands. CpG islands are present in the promotor regions of most human genes (60%). Methylation patterns play a role in cellular differentiation and tissue specific gene expression already during early development (41-43). The methylation pattern is continuously modified during mitosis and as part of DNA repair mechanisms and correct patterns needs to be maintained throughout life to grant cellular function (44, 45). Figure 4 visualizes the circular pathway of DNA methylation, hydroxylation and de-methylation and shows the mode of action of involved proteins and metabolites.

**Figure 4.**
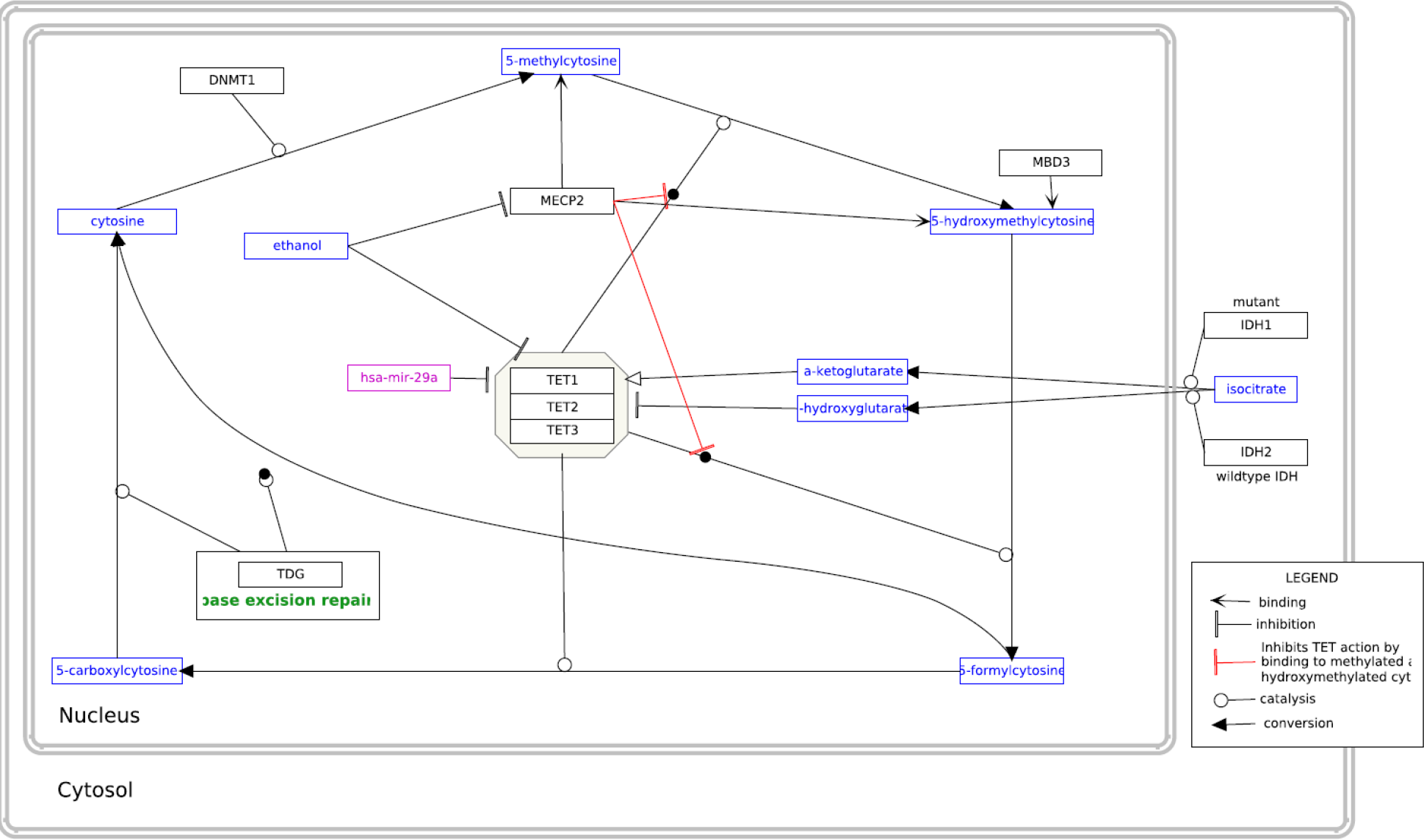
Pathway of cytosine methylation. The biological process is available atwikipathways.org/instance/WP3585.

MECP2 recognizes and binds specifically to 5MeCyt present in DNA. After binding it attracts co-repressor complexes containing SIN3A, NCOR and SMRT. This co-repressor complex finally recruits histone deacetylase (HDAC) (19, 30, 36, 46) (Figure 2, see MECP2 – HDAC complex). The complex acts by removing certain acetyl groups from histone proteins leading to chromatin condensation around methylated DNA (47, 48). Cell culture experiments with an inhibitor of HDAC resulted in the same phenotype as MECP2-KO cells (49). The NCOR-SMRT interaction domain (NID) of MECP2 is located within the TRD domain and the association with NCOR-SMRT is responsible for transcription repression activity of MECP2.

MECP2 was generally considered as a transcription inhibitor but recent research found also a conditional transcription activation function. Skene et al. identified MECP2, because of its mere amount, being equal to number of histone octamers or methylated DNA sites, as a global damper of gene transcription in mature mouse brain cells (32). During MECP2 absence H3 (histone family 3) acetylation levels were globally elevated and H1 (histone family 1) levels doubled suggesting MECP2 alters global chromatin state towards condensation and represses transcription. Li et al. on the other hand found global transcription activation by MECP2 in human embryonic stem cells which underwent differentiation to neuronal cells but repression by MECP2 in mature neuronal cells indicating that MECP2 can be both, an activator and repressor (50). The activation mechanism is explained as follows: MECP2 recruits CREB1 as a cofactor to target gene promotors (51) (Figure 2, Activation of transcription). MECP2 binding to 5OHMeCyt was even interpreted as a marker of active genes in neurons (51). MECP2 was also found to form a TET1 containing complex which leads to 5MeCyt hydroxylation and further to demethylation of DNA, enabling transcription (52). This mechanism was found to activate expression of downstream genes, namely BDNF, SST, OPRK1, GAMT, GPRIN1, and CREB1 (18) and is contradictory to other findings which describe MECP2 to block DNA demethylation by TET complex (Figure 4) (53).

MECP2 is additionally responsible for neuronal activity triggered transcription. Neuronal membrane depolarization and Ca^2+^ influx leads to phosphorylation of MECP2 which makes it detach from DNA, allowing decondensation and transcription (Figure 2) (54-58). Specific blocking of MECP2 phosphorylation sites led to RTT like symptoms (59).

## 6 Mutations of *MECP2* leading to RTT

At the moment, several hundred different mutations have been reported leading to RTT by loss or impaired function of MECP2 protein due to truncation, abnormal folding, or binding instability (15). This contributes to the variety of RTT phenotype and symptom severity.

The effects of total absence of MECP2 protein were investigated in several model systems. Deletion of *MECP2* from glial cells had only mild phenotypic consequences (60). In a mouse model specific deletion of *MECP2* in the forebrain caused behavioral abnormalities, limb clasping, impaired motor coordination, anxiety, and abnormal social behavior but not locomotor activity or changes in fear conditioning (61). In another mouse model, silencing MECP2 in GABAergic neurons led to severe RTT like phenotype (62).

67% of all *MECP2* mutations found in humans of are in eight hot spots: R106 (corresponding RS number from dbSNP: rs28934907), R133 (rs28934904), T158 (rs28934906), R168 (rs61748427), R255 (rs61749721), R270 (rs61750240), R294 (rs61751362) and R306 (rs28935468). Most of the mutations which cause RTT occur in the MDB region of *MECP2* (59). A 100fold reduction of binding affinity of MECP2 to methylated DNA is documented for the mutations R106W (rs28934907), R133C (rs28934904), and F155S (rs28934905) and binding affinity reduction of about 2fold was found in mutation T158M (rs28934906) (34).

## 7 The most important metabolites, genes and pathways affected by MECP2 in RTT

The examination and investigation of RTT females (and model systems) revealed that an impaired MECP2 influences biological pathways on many levels. Several genes have been found to be increased or decreased in expression, levels of various metabolites are changed and several biological pathways were found to be typically affected. In this chapter the main metabolites, genes and pathways which are influenced or changed by RTT are summarized. For getting these results, samples from human RTT females were often used but many results come from studies with Mecp2^-/y^ mice (e.g. the Bird model (63)). These mice do not express Mecp2 at all and they display the same symptoms as humans, such as normal development until about 6 weeks, regression, reduced movement, clumsy gait, irregular breathing, and the mice have a reduced life span of about 3 months. Postmortem analysis revealed reduced brain and neuronal cell size which is similar to observations in humans. Other mouse strains or *in vitro* models with mutated MECP2, reduced or overexpressed MECP2 levels are also commonly used to study RTT. Recently, researchers started to use iPSCs (induced pluripotent stem cells) from human or murine origin (64).

### Metabolites

RTT syndrome was originally named cerebroatropic hyperammonemia, indicating that one of the first measurable findings was an increased level of ammonium in blood (3). Early autopsies revealed reduced levels of catecholamines, namely dopamine, serotonin and norepinephrine while markers for bioaminergic metabolism in general were higher (65, 66) (summary of metabolites in Table 3 and see also the list of metabolites on wikipathways.org/instance/WP3584). This was confirmed by a study of Panayotiset al. (67) who revealed time-dependent levels of dopamine, norepinephrine, serotonin, and their catabolites in the brain tissues of Mecp2^-/y^ mice. Viemari (68) et al. showed that Mecp2^-/y^ mice have a deficiency in norepinephrine and serotonin content in the medulla and a drastic reduction of medullary TH (catecholamine producing) neurons indicating that dysfunctional neuronal development may be the reason for decreased catecholamine levels. The phosphorylation of MECP2 is triggered by neuronal activity which is caused by release of neurotransmitters (Figure 2). Hutchinson et al. (69) demonstrated that phosphorylation of MECP2 is dependent on dopamine, serotonin and norepinephrine activated pathways (Figure 2, MECP2 phosphorylation).

Another metabolite shown to be present in high levels in RTT females is glutamate. Increased glutamate production (or decreased glutamate consumption) may lead to over excitation of glutamatergic neurons and can trigger increased uptake and conversion of glutamate to glutamine. Over excitation feedback may cause the downregulation of the glutamate receptor which was observed before (70). Moreover, severe downregulation in gene expression for the glutamate D1 receptor GRID1 (GluD1) was found (Figure 2). This receptor links post and presynaptic compartments and influences not only synapsis function but also neuronal differentiation (71). Glutamate levels during sleep-wake cycle were also affected in Mecp2 deficient mice (72). Additionally, BDNF, which is directly regulated by MECP2, is also known to regulate GRID1 expression (73). Glutamate disposal is energy consumptive which may help explain the increased level of energy metabolism found in brains of Mecp2^-/y^ mice and RTT females in neuroimaging studies (74).

A metabolomics investigation found changed phospholipid profiles in Mecp2^-/y^ mice (70). Phospholipid metabolism is directly associated with cell growth since it provides membrane material for inner and outer hull structure. Still, it remains unclear whether this is the cause or just one of the consequences of reduced neuronal cell size and network connectivity of RTT. But as MECP2 basic function is global transcription dampener in combination with activity dependent activation of transcription this suggests that reduced membrane material production is a consequence of lack of activity specific transcription activation.

**Table 3:** Metabolites in RTT

| Changes in RTT | Metabolites |
| --- | --- |
| Increased levels (65-67) | Catabolites of catecholamine metabolism, glutamate |
| Decreased levels (65-68) | Dopamine, norepinephrine (noradrenaline), serotonin, melatonin, myo-inositol, phospholipids, GABA |

### Genes/gene products

MECP2 is a global transcription and translation influencing factor but there are also single specific genes which are found to be up- or downregulated in the absence of functional MECP2. The expression of the MECP2 target genes is affected in human, mouse and *in vitro* models. In the absence of functional MECP2 the expression of <u>BDNF, FKBP5, IGF2, DLX5, DLX6, SGK1, GAMT, FXYD1</u> and <u>MPP1</u> is upregulated whereas the expression of <u>UBE3A</u> (15) and <u>GRID1</u> (71) are downregulated (Table 4 and see also the list on wikipathways.org/instance/WP3584). BDNF, which shows in all investigated models the most consistent effect, is a MECP2 regulated protein and it is necessary for neuronal development and function, too (Table 4). <u>BioGRID</u> is a biological repository for interaction data. In this repository, there are 156 unique interactors for (human) MECP2 registered and 11 for BDNF. The current complete list can be found online using these links for <u>MECP2</u> and <u>BDNF</u>.

**Table 4:**
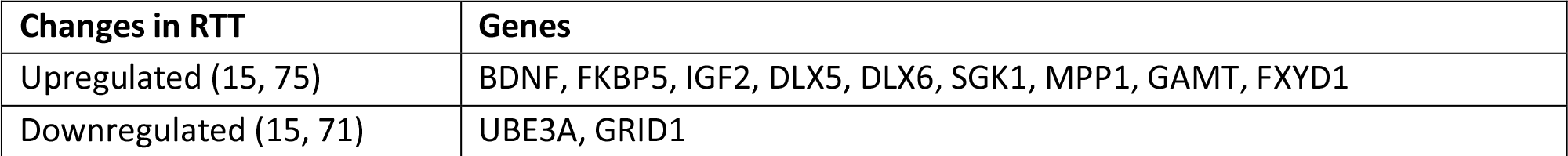
Genes, which are up- or downregulated in human or model system without functional MECP2

### Pathways

Bedogni et al. mentioned the difficulty to identify unique target pathways of MECP2 because MECP2 is both a repressor and an activator of transcription and the balancing and timing of transcription levels seems to contribute more to disorder development than activation of single pathways (59). Several transcriptomics studies using samples from RTT females, mouse and *in vitro* model systems showed about 60 significantly enriched pathways, for example inflammation, MAPK signaling, ERBB signaling, neurotropin intracellular cascade, sterol biosynthesis (76), cholesterol metabolism (76), cytoskeleton formation, and apoptosis (59).

While animal and *in vitro* models often use *MECP2* – KO models (e.g. Mecp2^-/y^ mouse) human RTT patient derived samples often have a residual MECP2 activity due to the various mutations which impair the function rather than inhibiting the expression completely. Tanaka et al. (64) investigated gene expression profile of iPSC lines derived from six different RTT females with different *MECP2* mutations. They created and compared expression profiles of two iPSCs cell lines derived from each of the patients, one with the X chromosome active that has the wild type *MECP2* and the other one with the chromosome with the mutated gene. The differently expressed genes and altered pathways are very different for each patient. This is an interesting result, as there are so many different *MECP2* mutations which lead to RTT. A specific mutation affects protein function differently and may trigger different pathways leading to different phenotypes. Linking genetic data to molecular analysis (transcriptome, metabolome) and phenotype will be a future challenge to elucidate the pathways of RTT.

## 8 MECP2-related pathways involved in RTT phenotypes

In this chapter we discuss how mutated *MECP2* leads to failure in neuronal synapsis formation and function which are one of the major causes of RTT phenotype. MECP2 acts in a biological network of constant interaction by regulating and being regulated. A phenotype is the final result of this complex network interaction.

According to Lyst and Bird (8) *MECP2* mutations cause RTT in two ways: 1. co-repressor recruitment, and 2. chromatin compaction, which are both basic molecular functions. Skene and Bedogni specified this assumption of MECP2 function as a global dampener of transcription in neurons plus the activity dependent transcription activation which leads to proper synapsis function and development (32, 59).

The differences in global gene expression of RTT and wild-type control groups are not substantial-neither in fold change nor number of genes differently expressed – indicating that more subtle dysregulation events in several pathways are responsible for RTT (51, 77-80). MECP2 is a global repressor of transcription (32), a global activator of gene translation (50) and it reacts on neuronal pathway signals which lead to phosphorylation of MECP2 and detachment from DNA. Therefore, MECP2 dampens neuronal transcription globally and allows activity related responses which seem to be necessary for learning activity and specific synapsis formation (81). Bedogni et al. found a direct connection between MECP2 and pathways leading to cytoskeleton re-formation (59) indicating structural changes leading towards neuronal network formation. Li et al. found in human embryonic stem cells, which are developing from stem cells to neuronal precursor cells to neurons, that in a premature state, MECP2 acts as an activator of transcription while transcription repressor activity was only found in mature neurons (50). The levels of MECP2 start to increase postnatally and the protein is quite abundant in mature nervous systems (32, 82). The expression of MECP2 is not uniform in different neuronal cell populations (83), parts of the brain and changes with age (84). Mouse Mecp2-KO neuronal precursor cells are not different from wildtype ones, they change only during maturation (82). Together with the observation that symptoms of RTT do not appear before about month 6, this led to the assumption that MECP2 has less to do with neurogenesis but more with neuronal function and maintenance, and synapsis formation and function. This may explain why the RTT phenotype becomes visible only at the quite late age of 6-18 months. In RTT females brains a decreased number of synapsis was found (85-87). Synaptogenesis again is mostly observed in the period of RTT symptom development (month 6 – 18) which may explain the development of learning disability. In MECP2 null mouse model reduced neuronal differentiation (88) and synaptic deficits (81) were observed. Mice studies and postmortem brains of RTT females reveal alterations in neuron structures which may be due to decreased dendritic complexity because of an immature synaptic spine morphology leading to malfunction of synaptic development and plasticity (89-91). Changed neuronal tubulin expression was found directly in the brain tissue of RTT and Angelman syndrome patients (92). Dysfunctional MECP2 led also to changes in synaptic transmission, short and long-term synaptic plasticity, deficits in short and long term potentiation (LTP and LTD) in mice (93).

The abnormal levels of neurotransmitters and differently expressed neurotransmitter receptors lead to an imbalance between excitatory and inhibitory neuronal activity (namely imbalance of GABAergic, glutamatergic and dopaminergic neuronal pathways). Such abnormal ratio of excitation/inhibition in brain activity was also found in autistic patients before (94-96) and it is a known effect in Parkinson’s disease which shares the motoric disabilities with RTT (97). RTT models using murine and human induced pluripotent stem cells showed some RTT features and these symptoms were documented for MECP2 overexpression models, too, leading again to the assumption that MECP2 function is dose dependent (16).

In summary, MECP2 affects epigenetic regulation of gene expression, which changes neurobiological activity, network formation and function, which causes the major phenotype. Although these processes could indeed explain many neuronal function related symptoms of RTT there is still lack of evidence for other phenotypes, especially those which occur in many but not in all RTT females. 1) Breathing patterns: breathing is regulated by brain stem function which gets its signals from receptors for blood pH, carbon dioxide and (to a lower amount) oxygen levels. Abnormal RTT breathing patterns could be caused by neuronal dysfunction of the brain stem or neuronal pathways but metabolic dysregulation involving abnormal creatin levels due to GAMT over/under expression is also an influencing factor (4). GAMT is one of the MECP2 downstream activated genes. 2) Cardiac abnormalities: heart beat is regulated by central nervous system but has its own nerve knots for signal production, too. Specialized neurons in the heart ensure proper electric signal transition. These might be affected by RTT directly, by brain stem function, or both. RTT patients are indeed known for higher incidence of heart problems. Furthermore, vascular dysfunctions have been found which are directly related to MECP2 dysfunction although the molecular pathway between MECP2 and effector genes is not yet elucidated (98). 3) Digestion and nutrient uptake problems: Stomach and intestines are covered with a complex network of nerve cells. Dysfunctional nerve cells may lead to constipation and malabsorption of nutrients, e.g. Vitamin D but there may also be other nutrient processing pathways involved. 4) Tissue specific effects: Tissue specific neuronal cells are differently influenced by MECP2 mutation, e.g. olfactory sensory vs. visual system (Table 5). Recently it was found that MECP2 mutations contribute to hypersensitivity of mechanoreceptors (99) which aligns with the observation of clinicians and caregivers. It is currently unknown in which neuronal cell subpopulations MECP2 is more or less necessary for normal function although there are indications that there are differences (83).

**Table 5a:**
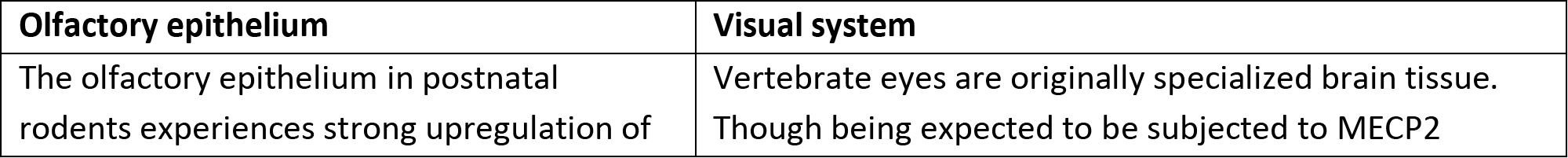
Olfactory vs. visual system: tissue specific MECP2 influence

**Table 5b:**
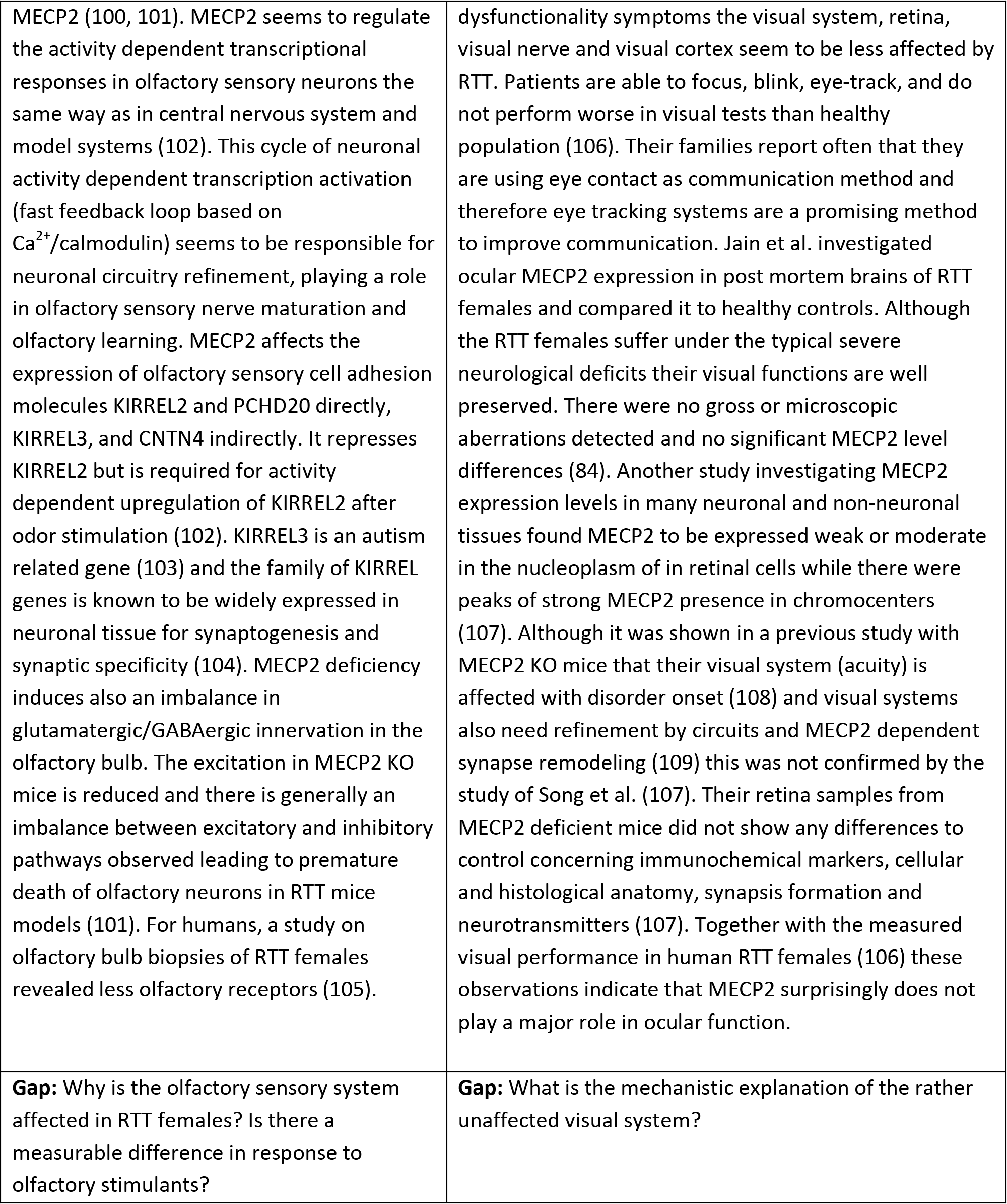

To understand these processes integration of different levels of biological knowledge and research results is necessary and interactive biological pathways (such as wikipathways.org/instance/WP3584) help to organize, analyze, and visualize existing knowledge. To integrate the information the exact mutation of MECP2 gene needs to be combined with molecular data (e.g. gene expression data, metabolomics) and a detailed description of the RTT females’ phenotype (including clinical laboratory measurements). This process will increase the understanding of the underlying pathways of the variety of RTT phenotypes. Gathering this knowledge and bringing it properly together needs collaboration of biomedical and bioinformatics researchers, physicians and patients (110). Furthermore, knowing the essential pathways and their components which contribute to a certain phenotype or symptom may lead to the discovery of drug targets. These are not likely to be able to cure RTT itself but may help to reduce the severity of specific symptoms.

## 9 Conclusion

The present review summarizes the current knowledge about MECP2 structure and function, how it influences levels of metabolites, gene expression and biological pathways, and tries to bridge the different types of data available to explain the development of typical RTT phenotype by visualizing in form of a pathway (Figure 2 and online wikipathways.org/instance/WP3584). Although the dysregulation events in neuronal cells can be already be explained quite well, the mechanistic explanation of several additional symptoms is still missing. Integrating the knowledge about the individual mutation, molecular data and phenotype information will help to find biological pathways and therefore explanations for these symptoms. Finding the right target genes, proteins, or metabolites can build a bridge between genotype and phenotype, and possibly to drug targets.

## 10 Acknowledgements

This work was funded by Stichting Terre – The Dutch Rett Syndrome Foundation.

### 11 Financial Disclosure

All authors report no biomedical financial interests or potential conflicts of interest.

